# Population structure of eulachon *Thaleichthys pacificus* from Northern California to Alaska using single nucleotide polymorphisms from direct amplicon sequencing

**DOI:** 10.1101/2020.05.31.126268

**Authors:** Ben J. G. Sutherland, John Candy, Kayla Mohns, Olivia Cornies, Kim Jonsen, Khai Le, Richard G. Gustafson, Krista M. Nichols, Terry D. Beacham

## Abstract

Eulachon *Thaleichthys pacificus*, a culturally and ecologically important anadromous smelt (Family Osmeridae), ranges from Northern California to the southeast Bering Sea. In recent decades, some populations have experienced declines. Here we use a contig-level genome assembly combined with previously published RADseq-derived markers to construct an amplicon panel for eulachon. Using this panel, we develop a filtered genetic baseline of 521 variant loci genotyped in 1,989 individuals from 14 populations ranging from Northern California through Central Alaska. Consistent with prior genetic studies, the strongest separation occurs among three main regions: from Northern California up to and including the Fraser River; north of the Fraser River to southeast Alaska; and within the Gulf of Alaska. Separating the Fraser River from southern US populations, and refining additional substructure within the central coast may be possible in mixed-stock analysis; this will be addressed in future work. The amplicon panel outperformed the previous microsatellite panel, and thus will be used in future mixed-stock analyses of eulachon in order to provide new insights for management and conservation of eulachon.

## INTRODUCTION

Eulachon *Thaleichthys pacificus* is a culturally and ecologically important anadromous smelt (Family Osmeridae) distributed in North America from Northern California to the southeast Bering Sea (Hay and McCarter 2000). Historically, approximately 95 rivers were considered to have spawning populations along the Northwest Pacific Coast, with large spawning populations in the Columbia River (USA) and the Fraser River (Canada) (COSEWIC 2011; Moody and Pitcher 2010). Eulachon are an important prey species for birds, marine mammals, and fishes, in part due to a high energetic benefit to cost ratio during foraging (Marston et al. 2002), as well as due to their returning to spawn at the end of winter and early spring when other prey species are scarce (Moody and Pitcher 2010). Eulachon are highly important to First Nations and American Indian indigenous peoples for both cultural and nutritional purposes, for example through the preparation and use of the rendered oil from adult eulachon, commonly known as “grease” (Moody and Pitcher 2010). Conservation and management of this species are therefore important goals in both Canada (DFO 2020) and the US (NMFS 2017).

Declines in eulachon populations have been reported coastwide since the mid 1990s, although some rivers maintain healthy returns including rivers in central Alaska (Ormseth 2018) and northern British Columbia (BC) (Hay and McCarter 2000). Adjacent regions both in southeast Alaska (SEAK) and Canada have declined in recent years (COSEWIC 2011). Although biomass has increased overall in Alaska eulachon populations, some specific runs in the area have had large reductions in returns (Flannery et al. 2013; Ormseth et al. 2008). In a meta-analysis by Moody and Pitcher (2010), factors were identified as associated with declines including bycatch in shrimp and hake fisheries, seal and sea lion predation, and increasing sea surface temperature. In Canada, no single threat could be identified for declines in abundance, although “mortality associated with coastwide changes in climate, fishing (direct and bycatch) and marine predation were considered to be greater threats… than changes in habitat or predation within spawning rivers” (DFO 2015; Schweigert et al. 2012). Forage fish population fluctuations are also dependent on environmental conditions, serving as the main link between climatic effects on primary producers and higher trophic levels (Guénette et al. 2014; Pikitch et al. 2014). In general, these fluctuations in forage fish populations are considered to be exacerbated by other factors, such as commercial fishing (Essington et al. 2015b), although the extent of this exacerbation is debated (Essington et al. 2015a; Szuwalski and Hilborn 2015).

Within Canada, the Committee on the Status of Endangered Wildlife in Canada (COSEWIC) considers eulachon to be in three Designatable Units (DUs): the Fraser River; the Central Pacific Coast; and the Nass/Skeena River (COSEWIC 2011). Both the Fraser River and the Central Pacific Coast DUs were assessed as endangered in 2011 (COSEWIC 2011), whereas the Nass/Skeena DU has been assessed as Special Concern (COSEWIC 2013). The Fraser River DU and the Central Pacific Coast DU remain under consideration for listing as endangered under Canada’s Species at Risk Act (SARA). Fisheries limitations and bycatch rules in Canada are outlined in Fisheries Management Plans (e.g., DFO 2020). Within the contiguous US, eulachon are designated as threatened under the Endangered Species Act within the southern Distinct Population Segment (DPS), ranging from northern California to the Skeena River, Canada (Gustafson et al. 2012; NMFS 2010). Improved characterization of population structure and the development of high-throughput methods to genotype unknown origin eulachon will help to better understand and manage this species, and will be useful in determining causes of declines.

In general, marine species with large population sizes are expected to have low population differentiation due to several factors including low effect of drift and high migration rate (Gagnaire et al. 2015), although this is not always the case, with molecular genetics now providing new resolution to previously considered homogenous stocks (Hauser and Carvalho 2008). Eulachon are less differentiated among populations than are other anadromous species such as salmonids (Candy et al. 2015), but are more differentiated than many marine species (Hauser and Carvalho 2008; Waples 1998). This may be due to a shorter in-river residence time, rapid downriver transport after hatch (Beacham et al. 2005; COSEWIC 2011; McLean and Taylor 2001), and thus less opportunity for imprinting, which may result in lower natal stream fidelity (Hay and McCarter 2000). Prior studies in eulachon using mitochondrial DNA markers were not able to identify population structure (McLean et al. 1999), but highly polymorphic microsatellite loci (Kaukinen et al. 2004) provided more resolution to define management units (Beacham et al. 2005). In Alaska, analyses with these same microsatellite loci identified two large population groupings, one to the north encompassing Cook Inlet, Prince William Sound, and Yakutat Forelands and one in the south from populations within the Alexander Archipelago south to Behm Canal (Flannery et al. 2013). This regional separation follows a hierarchical island model (Slatkin and Voelm 1991) and did not show isolation-by-distance within each region. The division of the regions and the lack of IBD within region may be in part due to larval dispersal associated with the counter-clockwise Alaska gyre dispersing larvae from the Yakutat Forelands to the east; the southern populations within the Alexander Archipelago would not experience the same transport. Restriction-site associated DNA sequencing (RADseq) was applied to eulachon collections from 12 populations spanning from the Columbia River to the Northern Gulf of Alaska (Candy *et al*. 2015). Although sample sizes for the RADseq project were smaller than those previously analyzed by microsatellites, this study provided strong evidence for three genetic clusters (i.e., Northern Gulf of Alaska, Southeast Alaska-BC, and Fraser-Columbia), and markers identified as putative outliers or having high F_ST_ were identified to be used for future studies. High F_ST_ markers can provide increased discrimination power for genetic stock identification (GSI; Ackerman et al. 2011; Hess et al. 2011). The combination of highly resolving markers and high sample sizes per population provides the best power for resolving a species with such low structure. However, such studies would benefit from having multi-year collections to ensure allele frequencies are stable over time (i.e., multiple brood cycles) and not highly impacted by sweepstakes reproductive success (Hedgecock and Pudovkin 2011) and annual variation within rivers (Waples 1998).

Assigning unknown origin fish by GSI and mixed-stock analysis (MSA) to population or region of origin is useful for interpreting at-sea ecology during surveys or bycatch interceptions, as little is known as to how eulachon mix among populations at-sea, or where populations are during different seasons. Knowing where specific DUs exist and are intercepted as bycatch may help reduce impacts on at-risk populations. Eulachon MSA has been conducted on eulachon bycatch from fisheries off the west coast of British Columbia since the early 2000s using the microsatellite panel (Beacham et al. 2005). Increased knowledge regarding eulachon at-sea ecology and behaviour, as well as the ability to identify natal origins of intercepted fish have both been highlighted as priority research needs in Canada (Schweigert et al. 2012), on the US West Coast (NMFS 2017), and in Alaska (Ormseth et al. 2008). The microsatellite panel used for MSA identified that eulachon bycatch from a Chatham Sound (North Coast of BC) shrimp trawl fishery in 2001 were predominantly of central mainland and Nass River origin, whereas eulachon caught in a research survey in Queen Charlotte Sound (Central Coast of BC) were from various regions coastwide, and those caught in a survey along West Coast Vancouver Island were predominantly of Columbia and Fraser origin (Beacham et al. 2005). Tests of samples of spawning eulachon in river mouths (i.e., genetic baseline collections) within central and southeast Alaska also have demonstrated the potential for assignment within the northern and southern groupings (Flannery et al. 2013). Although ultimately the resolution ability for GSI depends on the level of differentiation that exists within the species, advances in genotyping technology makes it possible to develop an improved baseline with highly resolving markers, large sample sizes, and multiple sampling years, providing more accurate estimates for MSA and GSI.

Low-density (i.e., ∼500 markers) single nucleotide polymorphism (SNP) marker panels genotyped using high-throughput sequencing of amplicons (also known as GTseq; Campbell et al. 2015) are routinely applied for GSI and parentage-based tagging (PBT) in order to genotype thousands of individuals at low costs, particularly in the salmonid fishes. For example, genetic baselines of Coho Salmon *Oncorhynchus kisutch* are currently being used for MSA in BC fisheries in combination with PBT to assign unknown origin individuals back to genotyped parents from hatchery broodstock (Beacham et al. 2019). GTseq is a cost-effective solution for questions that do not require high-density marker panels, but that need to be applied to thousands of individuals (Campbell et al. 2015; Meek and Larson 2019). Several aspects of SNPs are advantageous compared to microsatellite markers, which include automation, cost, portability of methods, and scalability (Hauser et al. 2011). Further, due to the fewer number of alleles per locus compared with microsatellite loci, fewer individuals are needed in order to generate allele frequencies for baseline samples (Beacham et al. 2011).

The present study aimed to develop an approximately 500 marker SNP panel using a contig-level genome assembly (Sutherland et al. in prep) and the top discriminating markers from the previous coastwide RADseq study (Candy et al. 2015). The developed SNP panel was applied to tissue archives and new collections to generate a representative baseline encompassing 1,989 individuals from 14 populations with at least 35 individuals per population from Northern California to Central Alaska. This baseline allowed us to compare resolution and power against the microsatellite baseline, to estimate minimum required sample sizes per population, and to evaluate temporal variation within each river system that contained collections from multiple years. The overarching goal of the SNP panel and eulachon genetic baseline is to ultimately use the SNP baseline to improve our understanding of population delimitations and at-sea ecology of eulachon via GSI and MSA.

## MATERIALS AND METHODS

### Amplicon panel design

Markers were obtained from a RADseq study of eulachon populations (n = 12 populations) from the Columbia River (Washington, USA) to the Northern Gulf of Alaska (Candy et al. 2015). These 4,104 RAD loci were each 90 bp in length and were aligned to the eulachon reference genome (Sutherland et al. in prep) using BLAST (Altschul et al. 1990) with a minimum e-value cutoff of 1e-20. Only the loci that aligned to the reference genome two or fewer times were retained in order to reduce non-specific hits and markers present in repetitive regions of the genome. The allowance of two alignments was due to the potential for redundancy in the reference contig assembly, and further evaluation of each designed marker was conducted after the panel design (e.g., excess heterozygosity). The top hit as evaluated by bitscore, e-value, and percent identity per locus was used to retain a single representative hit per locus for downstream analysis.

A total of 200 bp in each direction flanking the first SNP in the marker was extracted from the reference genome using a custom pipeline (see Data Accessibility: *fasta_SNP_extract*). If the start or end of a contig was less than 200 bp from the SNP, then a 400 bp segment was obtained from the start or end of the contig, respectively. The segment of interest was extracted from the contig using *bedtools* (Quinlan and Hall 2010). The variant locus was marked with both alleles within the output amplicon fasta file. All putatively adaptive markers from Candy et al. (2015; n = 193) that passed quality filters were retained for the design (n = 181 putatively adaptive markers). Putatively neutral markers with the highest overall F_ST_ were added until a total of 600 markers were present in the panel (n = 419 neutral markers). The panel was then submitted for design through the AgriSeq panel design pipeline (Thermo Fisher).

### Sequencing and variant calling

Eulachon samples in the tissue archive at the Molecular Genetics Laboratory (MGL) at Pacific Biological Station (Fisheries and Oceans Canada) or provided by collaborators (Table 1; see *Acknowledgements*) were amplified using the designed panel as per manufacturers’ instructions (Thermo Fisher).

**Table 1.**
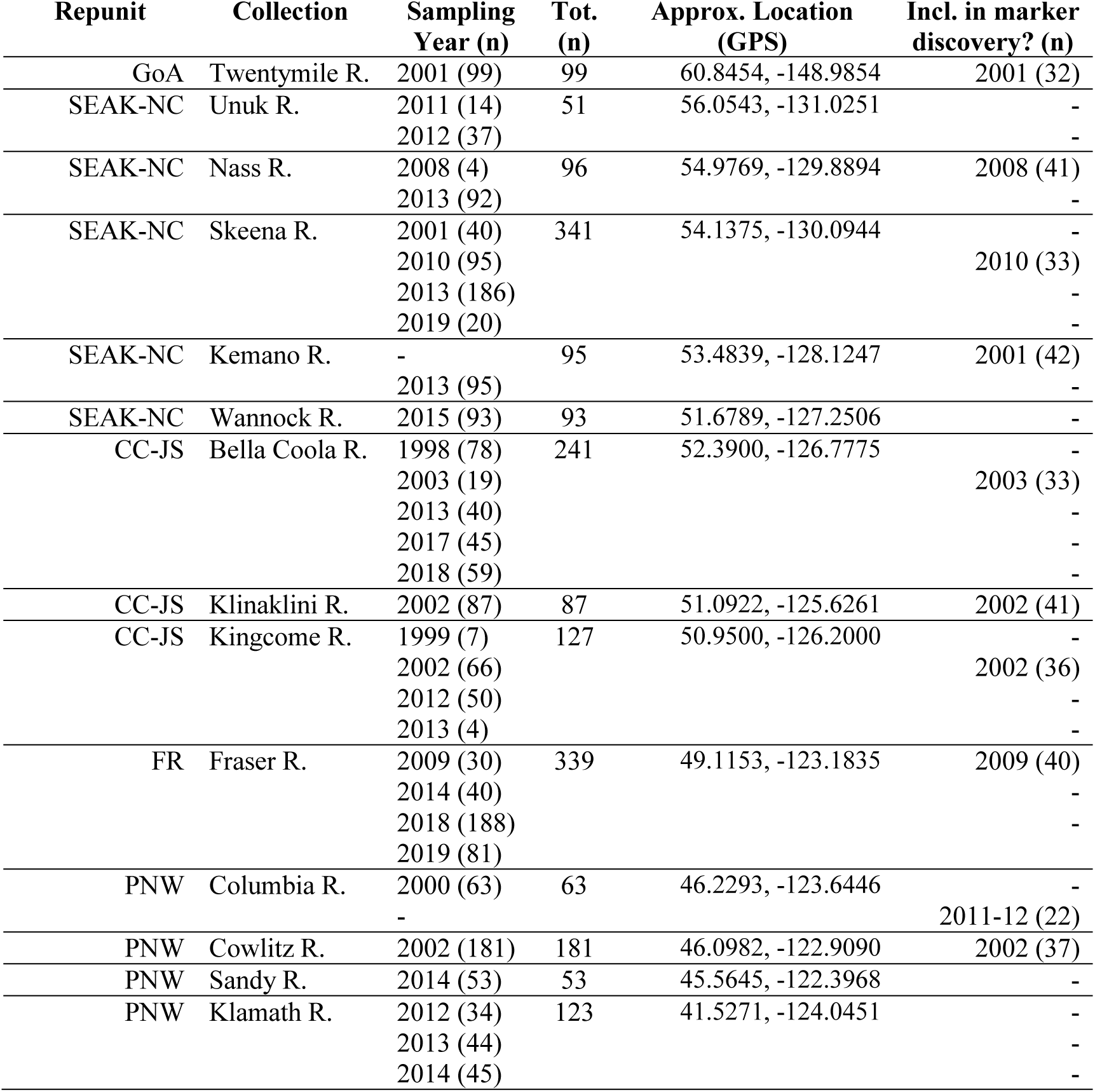
Populations and years included in the study retained after quality control of baseline shown within larger groupings (i.e., repunits) and approximate location (GPS). The number of samples from each year as well as the total number of samples for the collection are shown (n). Collections are also shown whether each was in the original marker discovery (i.e., Candy et al. 2015).

| Repunit | Collection | Sampling<br>Year (n) | Tot.<br>(n) | Approx. Location<br>(GPS) | Incl. in marker<br>discovery? (n) |
| --- | --- | --- | --- | --- | --- |
| GoA | Twentymile R. | 2001 (99) | 99 | 60.8454, -148.9854 | 2001 (32) |
| SEAK-NC | Unuk R. | 2011 (14) | 51 | 56.0543, -131.0251 | - |
|  |  | 2012 (37) |  |  | - |
| SEAK-NC | Nass R. | 2008 (4) | 96 | 54.9769, -129.8894 | 2008 (41) |
|  |  | 2013 (92) |  |  | - |
| SEAK-NC | Skeena R. | 2001 (40) | 341 | 54.1375, -130.0944 | - |
|  |  | 2010 (95) |  |  | 2010 (33) |
|  |  | 2013 (186) |  |  | - |
|  |  | 2019 (20) |  |  | - |
| SEAK-NC | Kemano R. | - | 95 | 53.4839, -128.1247 | 2001 (42) |
|  |  | 2013 (95) |  |  | - |
| SEAK-NC | Wannock R. | 2015 (93) | 93 | 51.6789, -127.2506 | - |
| CC-JS | Bella Coola R. | 1998 (78) | 241 | 52.3900, -126.7775 | - |
|  |  | 2003 (19) |  |  | 2003 (33) |
|  |  | 2013 (40) |  |  | - |
|  |  | 2017 (45) |  |  | - |
|  |  | 2018 (59) |  |  | - |
| CC-JS | Klinaklini R. | 2002 (87) | 87 | 51.0922, -125.6261 | 2002 (41) |
| CC-JS | Kingcome R. | 1999 (7) | 127 | 50.9500, -126.2000 | - |
|  |  | 2002 (66) |  |  | 2002 (36) |
|  |  | 2012 (50) |  |  | - |
|  |  | 2013 (4) |  |  | - |
| FR | Fraser R. | 2009 (30) | 339 | 49.1153, -123.1835 | 2009 (40) |
|  |  | 2014 (40) |  |  | - |
|  |  | 2018 (188) |  |  | - |
|  |  | 2019 (81) |  |  | - |
| PNW | Columbia R. | 2000 (63) | 63 | 46.2293, -123.6446 | - |
|  |  | - |  |  | 2011-12 (22) |
| PNW | Cowlitz R. | 2002 (181) | 181 | 46.0982, -122.9090 | 2002 (37) |
| PNW | Sandy R. | 2014 (53) | 53 | 45.5645, -122.3968 | - |
| PNW | Klamath R. | 2012 (34) | 123 | 41.5271, -124.0451 | - |
|  |  | 2013 (44) |  |  | - |
|  |  | 2014 (45) |  |  | - |

DNA was extracted from tissues using a variety of methods, including chelex extraction, a modified version of the chelex extraction (i.e. PBT chelex) that includes an additional overnight incubation (with Proteinase-K, chelex beads and UltraPure water) and no high temperature thermal cycling step, the Wizard SV 96 Genomic DNA Purification System (Promega), DNeasy (QIAGEN), and BioSprint (QIAGEN). Highest genotyping rates were observed using DNeasy or BioSprint methods, and so when possible, these were used for additional extractions.

Individuals were barcoded and amplified using the AgriSeq panel protocol and the eulachon v.1.0 primers (Additional File S1) as per manufacturers instructions, using the 768 barcodes available (Thermo Fisher), as previously described (Beacham et al. 2017). Individuals were multiplexed in batches of 768 and sequenced on each Ion Torrent PI chip (Thermo Fisher). Sequenced samples were de-multiplexed and variants called using a hotspots file (Additional File S2) in the Torrent Suite software (TS v.5.10.1; *variantCaller* v.5.6.0.4; Thermo Fisher). Called variants were then imported into the MGL genotype database management system.

### Data filtering

Data filtering was conducted sequentially, where first, samples were removed due to missing genotypes (i.e., retain individuals with less than 50% missing data, that is, being genotyped at >259 amplicons). The cutoff of >259 amplicons was chosen as 50% of the total ∼518 amplicons that typically made it through quality control filters (note: this number changed slightly depending on the samples included in the analysis). Second, populations with too few samples were removed (i.e., retain when population has ≥ 20 individuals) for initial evaluation of population structure. Third, amplicons with excess heterozygosity (≥ 0.5), or those missing in too many individuals (≥ 50% samples) were removed from the data. Markers were previously screened for deviations from Hardy-Weinberg equilibrium (Candy et al. 2015). The filtered baseline database was date-stamped and converted to genepop format for downstream genetic analyses.

Locations of baseline populations were plotted on a map in R (R Core Team 2020) using ggplot2 (Wickham 2016) and ggrepel (Slowikowski 2019) based on the GPS coordinates at the river mouth of baseline sites.

### Population differentiation analysis

The baseline genotypes for all individuals were read into R using adegenet (Jombart 2008) using a custom pipeline (see Data Accessibility; *simple_pop_stats*). Pairwise F_ST_(Weir and Cockerham 1984) including 95% confidence intervals was calculated using *pairwise*.*WCfst* and *boot*.*ppfst* for all populations or year-separated populations using hierfstat (Goudet 2005). A neighbour-joining tree using the *edwards*.*dist* distance metric (Cavalli-Sforza and Edwards 1967) was generated using the aboot function of *poppr* (Kamvar et al. 2014) with 10,000 bootstraps. This was exported in tree format and input to FigTree v1.4.4 for data visualization (Rambaut 2019). This was conducted for all populations with at least 20 individuals, then with all populations with at least 35 individuals. It was also conducted for all populations with at least 35 individuals separated by year.

Isolation-by-distance (IBD) was evaluated by finding an approximate distance between all recorded GPS coordinates for all collection sites using the *distm* function of geosphere (Hijmans 2019) and custom scripts (Data Accessibility; *simple_pop_stats*). Pairwise F_ST_ and pairwise physical distances (km) were compared to calculate a linear model best fit line to determine adjusted R^2^ values of how well the data fit the model. Within-region IBD was investigated for both northern and southern populations. IBD across regions was investigated using all populations with at least 20 individuals, then with all populations with at least 35 individuals to estimate the effect of population sample size on adherence to IBD.

To determine the proportion of variance captured within larger groupings observed in the dendrogram, within populations inside larger groupings (i.e., to determine the amount of variation among collections within a larger grouping), and the unexplained remaining variation that exists among samples, an Analysis of Molecular Variance (AMOVA) was calculated using the function *poppr*.*amova* of poppr that uses the *ade4* package (Dray and Dufour 2007) using default parameters (e.g. removing loci with more than 5% missing data).

### Multivariate statistics

Principal Components Analysis (PCA) was performed by first converting the genind to genlight using the *gi2gl* function of dartR (Gruber et al. 2018), then conducting a genlight PCA by the glPca function of adegenet, then plotting with ggplot2 using 95% confidence to draw ellipses around the samples from each grouping using the function *stat_ellipse* (Wickham 2016). Eigenvalues were plotted, three principal components were retained, and allele loadings for each PC as characterized by *loadingplot* in adegenet were plotted to identify top loading markers into PCs. Further, a Discriminant Analysis of Principal Components (DAPC) was performed using adegenet, retaining 10 PCs and one axis, and variance contributions were plotted. Top loading markers in the DAPC were characterized.

### Relatedness

Inter-individual relatedness within a population was calculated for all populations to compare relative relatedness values. A genlight object was created using dartR, then converted to Demerelate format (Kraemer and Gerlach 2017) in order to format using the ‘readgenotypedata’ function of related (Pew et al. 2015). Subsequently, the coancestry was calculated within related, implementing *coancestry* (Wang 2011) using the *ritland* (Ritland 1996) and *wang* (Wang 2002) metrics. Relatedness for these metrics was plotted in R for relatedness within a population.

### Microsatellite Data

Microsatellite data were obtained to compare with the baseline data genotyped by the SNP panel. Existing microsatellite data were obtained from the baseline collection database at MGL (Beacham et al. 2005) using Microsatellite Manager v.10.3 (Candy et al. 2002). The newly added population from California (Klamath River) was genotyped using the same genotyping methods as previously described (Beacham et al. 2005). F_ST_, IBD, and dendrograms were all calculated as described above.

## RESULTS

### Amplicon panel design

Of the total 4,104 single SNPs in RAD-tags from Candy et al. (2015), 3,957 (96%) were found to have at least one significant alignment against the eulachon reference genome (Sutherland et al. in prep). From these markers, 3,880 aligned with a single significant hit, 48 with two hits, and 29 with more than two hits. RAD loci with two or fewer hits were retained for further development (n = 3,928 RAD loci). These markers were further reduced to a total of 600 SNPs by preferentially selecting the putatively adaptive loci that passed alignment filters (n = 181 of 193) and high F_ST_ (n = 419) SNPs from Candy et al. (2015). Of these, six putative adaptive and 14 neutral SNPs submitted did not pass primer design, leaving a total of 580 pairs designed into the eulachon AgriSeq panel (v1.0; Thermo Fisher). This panel is comprised of primers optimized for Ion Torrent Proton technology, and has not been tested with other technology. Primer sequences are available in Additional File S1.

### Baseline population genotyping and quality control

Four sequencing chips (PI v3; Ion Torrent) were used for direct amplicon sequencing of baseline populations of up to 768 individuals per chip. Sequencing generated a total of 83.9 M, 58.5 M, 79.7 M, and 48.9 M reads within amplicons per chip for baseline samples, with an average number of reads per sample of 123 k (median = 102 k; standard deviation = 112 k), 77 k (med = 33 k; sd = 121 k), 105 k (med = 73 k; sd = 114 k), and 87 k (med = 65 k; sd = 83 k), respectively. After individual samples with high missing data were removed, a total of 19 populations with at least 20 individuals per population was retained (Table 1). The amplicons, not including primer sequence, were on average 168 bp (min = 84 bp; max = 188 bp). Markers were identified that had excess heterozygosity (Figure S1; n = 35 markers) or excess missing data across samples (n = 19). Removing these markers left a total of 526 amplicons. Five of these markers were monomorphic across the populations genotyped, leaving 521 remaining amplicons, with a single SNP designated per amplicon. This reduced set of markers is the quality controlled marker set (Additional File S2). After all of the quality control, on average there were 111 samples per population (sd = 98 samples).

To determine the appropriate minimum sample size threshold per population, all populations with at least 20 individuals were clustered into a dendrogram (Figure S2). The groupings of populations with 35 or fewer individuals (i.e., Bear River, Falls Creek, Kitimat River, Carroll Creek, and Elwha River) were consistently outside of the main clusters. Therefore, a threshold of 35 individuals per population was applied (see Figure 1) and any populations with fewer than 35 individuals were removed from the baseline. This resulted in a total of 1,989 individuals for 14 populations, with an average of 142 individuals per population (sd = 98; min = 51; max = 339).

**Figure 1.**
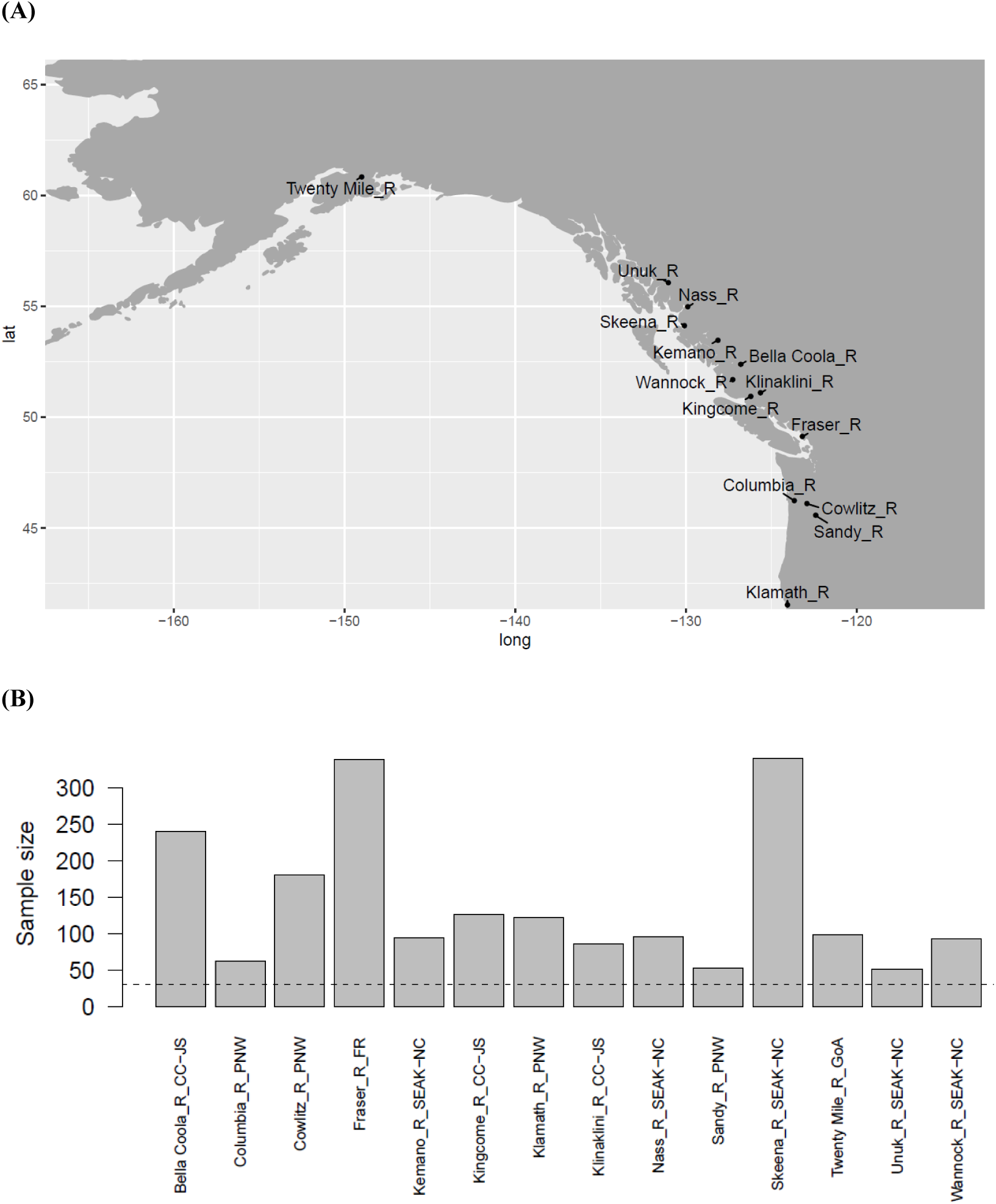
Locations (A) and sample sizes (B) for all collections with more than 35 individuals per collection included in the filtered amplicon baseline.

### Hierarchical population structure

The most divergent population in the dataset was Twentymile River (AK), a river at the upper end of Turnagain Arm in south-central Alaska (Figure 1). This was indicated by the relatively high genome-wide differentiation when compared to all other populations (mean F_ST_ = 0.0427; Table S1). For comparison, a population near the middle of the sampled range, Klinaklini River, compared with all other populations except Twentymile River indicates much less differentiation (mean F_ST_ = 0.0100). This distinctiveness of Twentymile River also can be observed in a Principal Components Analysis (PCA) along PC2 (Figure S3). PC2 was separated by numerous markers, although some contributed more substantially to the division (Figure S4A).

The second largest separation in the data separated populations from the Fraser River and south, grouping the Fraser, Columbia, and Klamath Rivers (Figure 2). This separation of the northern and southern clusters had high bootstrap support (> 99.99 %), and was also apparent along PC1 of the PCA (Figure S3). PC1 was also separated by numerous markers, although ∼8 markers showed a high contribution to this separation (Figure S4B). Within the southern grouping, there was some clustering of Columbia River populations together, but the Cowlitz River population, the most numerous of the Columbia River collections and entirely sourced from 2002 (Table 1), grouped into a cluster with Klamath River, and more broadly with the Fraser River (Figure 2), rather than with the other Columbia River populations (Columbia River, Sandy River). Cowlitz River and Klamath River are grouped closely together and in 87% of trees group together without the Fraser River. In general these populations were very similar (e.g., Fraser River vs. Columbia River F_ST_ = 0.0079, 95% confidence interval (CI): 0.0044-0.0130; Fraser River vs. Klamath River F_ST_ = 0.0021, 95% CI: 0.0012-0.0030; and Klamath River vs. Columbia River F_ST_ = 0.0091, 95% CI: 0.0051-0.0146; Table 2).

**Table 2.**
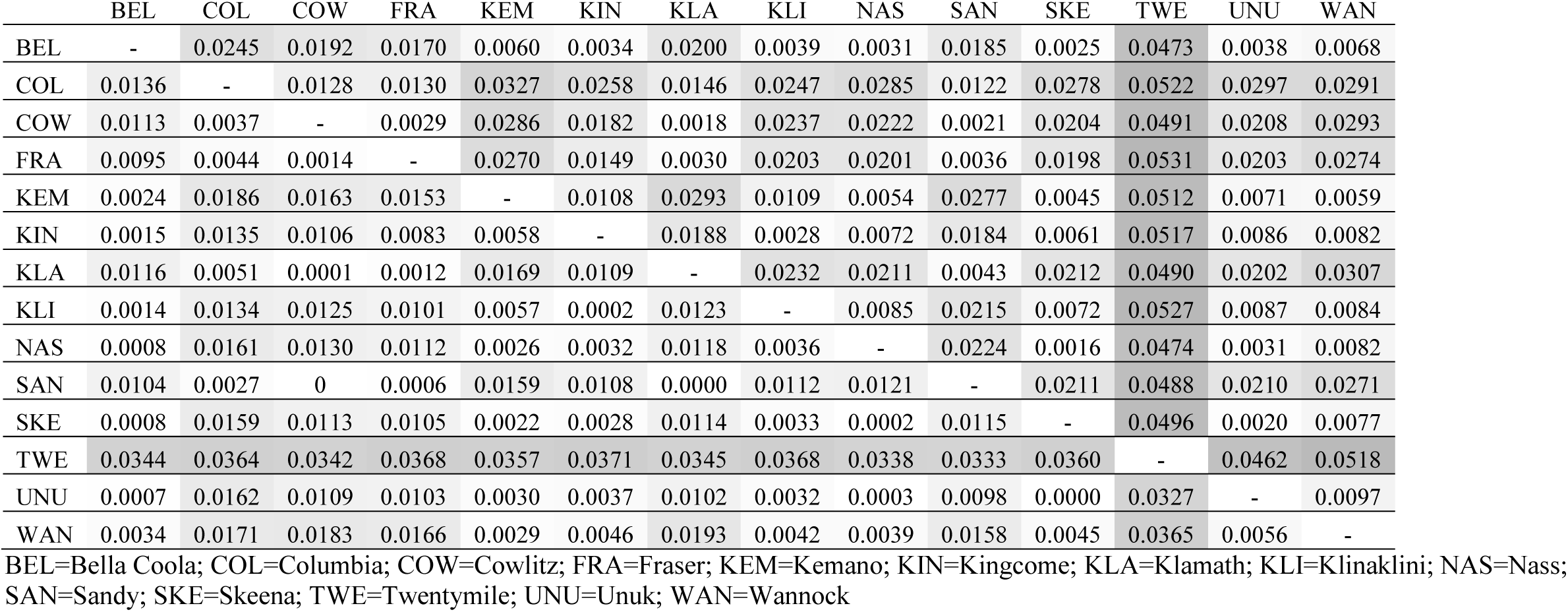
Pairwise genetic differentiation between populations using the SNP panel as shown by Weir-Cockerham F_ST_ 95% confidence limits (lower limits in the bottom half, upper limits in the upper half). Shading is used to show increasing values. Only populations with more than 35 individuals are shown. Negative values in the lower limit were replaced by zero and the comparison was considered to be not significantly different.

|  | BEL | COL | COW | FRA | KEM | KIN | KLA | KLI | NAS | SAN | SKE | TWE | UNU | WAN |
| --- | --- | --- | --- | --- | --- | --- | --- | --- | --- | --- | --- | --- | --- | --- |
| BEL | - | 0.0245 | 0.0192 | 0.0170 | 0.0060 | 0.0034 | 0.0200 | 0.0039 | 0.0031 | 0.0185 | 0.0025 | 0.0473 | 0.0038 | 0.0068 |
| COL | 0.0136 | - | 0.0128 | 0.0130 | 0.0327 | 0.0258 | 0.0146 | 0.0247 | 0.0285 | 0.0122 | 0.0278 | 0.0522 | 0.0297 | 0.0291 |
| COW | 0.0113 | 0.0037 | - | 0.0029 | 0.0286 | 0.0182 | 0.0018 | 0.0237 | 0.0222 | 0.0021 | 0.0204 | 0.0491 | 0.0208 | 0.0293 |
| FRA | 0.0095 | 0.0044 | 0.0014 | - | 0.0270 | 0.0149 | 0.0030 | 0.0203 | 0.0201 | 0.0036 | 0.0198 | 0.0531 | 0.0203 | 0.0274 |
| KEM | 0.0024 | 0.0186 | 0.0163 | 0.0153 | - | 0.0108 | 0.0293 | 0.0109 | 0.0054 | 0.0277 | 0.0045 | 0.0512 | 0.0071 | 0.0059 |
| KIN | 0.0015 | 0.0135 | 0.0106 | 0.0083 | 0.0058 | - | 0.0188 | 0.0028 | 0.0072 | 0.0184 | 0.0061 | 0.0517 | 0.0086 | 0.0082 |
| KLA | 0.0116 | 0.0051 | 0.0001 | 0.0012 | 0.0169 | 0.0109 | - | 0.0232 | 0.0211 | 0.0043 | 0.0212 | 0.0490 | 0.0202 | 0.0307 |
| KLI | 0.0014 | 0.0134 | 0.0125 | 0.0101 | 0.0057 | 0.0002 | 0.0123 | - | 0.0085 | 0.0215 | 0.0072 | 0.0527 | 0.0087 | 0.0084 |
| NAS | 0.0008 | 0.0161 | 0.0130 | 0.0112 | 0.0026 | 0.0032 | 0.0118 | 0.0036 | - | 0.0224 | 0.0016 | 0.0474 | 0.0031 | 0.0082 |
| SAN | 0.0104 | 0.0027 | 0 | 0.0006 | 0.0159 | 0.0108 | 0.0000 | 0.0112 | 0.0121 | - | 0.0211 | 0.0488 | 0.0210 | 0.0271 |
| SKE | 0.0008 | 0.0159 | 0.0113 | 0.0105 | 0.0022 | 0.0028 | 0.0114 | 0.0033 | 0.0002 | 0.0115 | - | 0.0496 | 0.0020 | 0.0077 |
| TWE | 0.0344 | 0.0364 | 0.0342 | 0.0368 | 0.0357 | 0.0371 | 0.0345 | 0.0368 | 0.0338 | 0.0333 | 0.0360 | - | 0.0462 | 0.0518 |
| UNU | 0.0007 | 0.0162 | 0.0109 | 0.0103 | 0.0030 | 0.0037 | 0.0102 | 0.0032 | 0.0003 | 0.0098 | 0.0000 | 0.0327 | - | 0.0097 |
| WAN | 0.0034 | 0.0171 | 0.0183 | 0.0166 | 0.0029 | 0.0046 | 0.0193 | 0.0042 | 0.0039 | 0.0158 | 0.0045 | 0.0365 | 0.0056 | - |

BEL=Bella Coola; COL=Columbia; COW=Cowlitz; FRA=Fraser; KEM=Kemano; KIN=Kingcome; KLA=Klamath; KLI=Klinaklini; NAS=Nass; SAN=Sandy; SKE=Skeena; TWE=Twentymile; UNU=Unuk; WAN=Wannock

**Figure 2.**
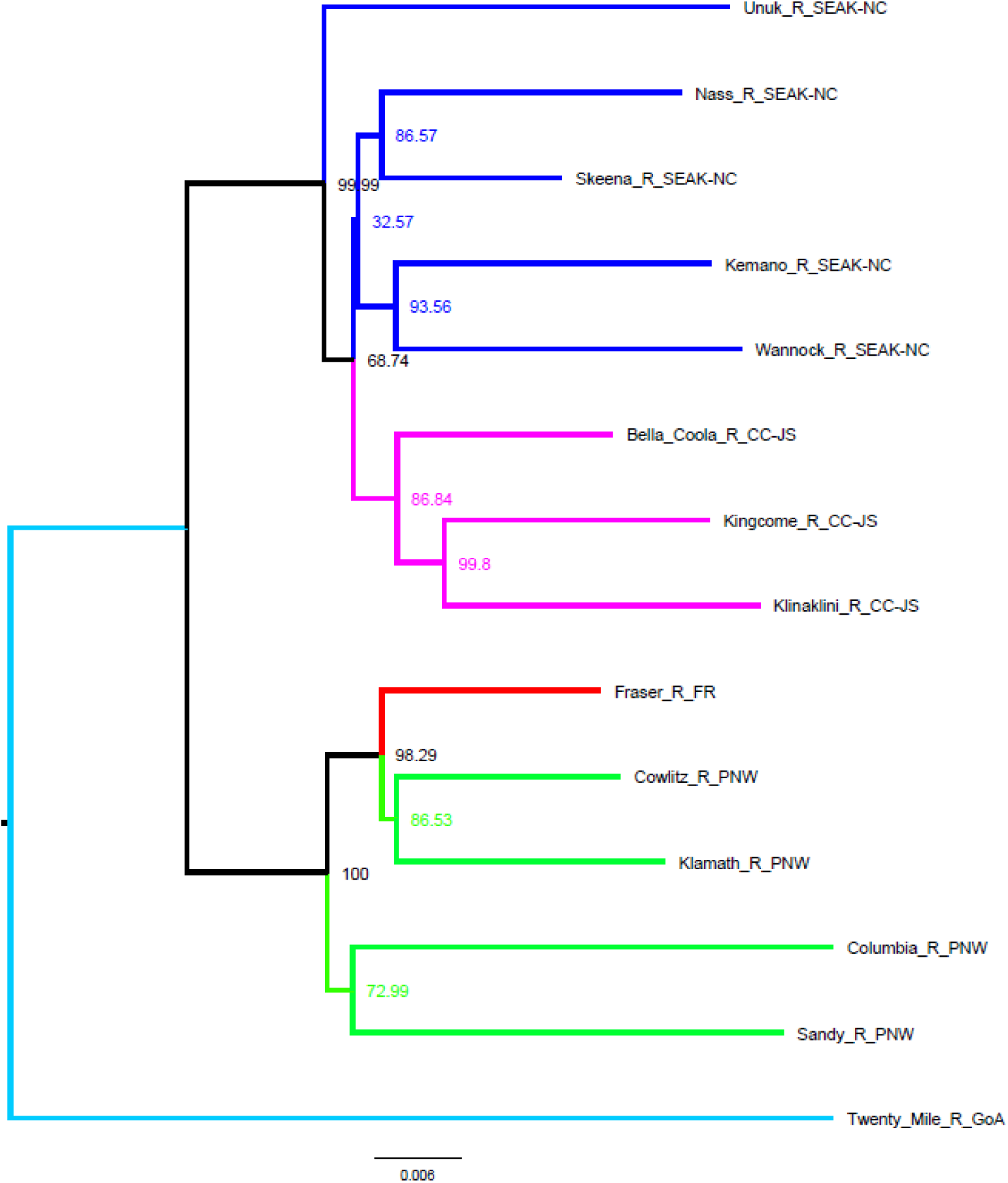
Genetic similarity among populations in the filtered amplicon baseline as shown within a neighbour-joining tree (Cavelli-Sforza and Edwards chord distance) rooted with Twentymile River. Branches are coloured by general grouping as shown in Table 1 (light blue = Gulf of Alaska; dark blue = Southeast Alaska, North Coast, and Central Coast; pink = additional Central Coast and Johnstone Strait populations; red = Fraser River; green = US southern populations).

Within the northern grouping, there was a strongly supported cluster including the populations in Johnstone Strait (Kingcome River and Klinaklini River; 99.8% bootstrap support) and more broadly with Bella Coola (86.84% bootstrap support; Figure 2). These populations had high genetic similarity with each other (mean F_ST_ = 0.0021; Table S1). Second, other Central Coast populations Kemano River and Wannock River were grouped together and were highly similar (93.56% bootstrap; F_ST_ = 0.0043, 95% CI: 0.0029-0.0059). Although Bella Coola River and Wannock River did not group in the same cluster as may be expected due to physical proximity, they still showed low differentiation (F_ST_ = 0.005, 95% CI: 0.0034-0.0068). North Coast populations Nass River and Skeena River were nearly indistinguishable (87% bootstrap; F_ST_ = 0.0009, 95% CI: 0.0002-0.0016). The Nass River was more differentiated from the Klinaklini River, for example (F_ST_ = 0.006, 95% CI: 0.0036-0.0085). The Transboundary Region’s Unuk River clustered outside of the North Coast and Central Coast groupings, but still within the larger northern grouping. Importantly, Bella Coola River also showed low differentiation from both the Skeena River, and the Unuk River (F_ST_ = 0.0016, 95% CI: 0.0008-0.0025; and F_ST_ = 0.0021, 95% CI: 0.0007-0.0038, respectively).

Although there are three clear groupings in the data (Gulf of Alaska, northern populations, southern populations), as expected from previous work (Candy et al. 2015), and initially appear to indicate evidence for Isolation-by-Distance (IBD) with a linear relationship between pairwise F_ST_ and physical distance (km; adjusted R^2^ = 0.71; Figure 3A), the observed IBD does not exist within regions, where the populations in the southern region and northern region do not individually show IBD (adj. R^2^ = 0.009 and 0.166, respectively; Figure 3C and 3D), and thus are more reflective of a hierarchical island model. Using five tentative reporting units (repunits; i.e., grouping of similar populations) as viewed in the dendrogram and shown by appended regional information in Figure 2, an analysis of molecular variation (AMOVA) was used to view the partitioning of variance within groupings. Although the majority of variation is among individuals within populations (96.34%), the between repunit variation was 2.13%, whereas the between samples within repunit was 1.53% (Table 3).

**Table 3.** Analysis of molecular variance (AMOVA) results showing sources of variation within the amplicon panel baseline, using the filtered baseline (i.e., greater than 35 individuals per collection, grouped by repunit as per Table 1).

| Source of variation | Degrees of freedom | Sums of squares | Variance components (sigma) | Percentage of variation |
| --- | --- | --- | --- | --- |
| Between repunit | 4 | 2268.21 | 1.0176 | 2.13 |
| Between samples within repunit | 9 | 1162.54 | 0.7301 | 1.53 |
| Within samples | 1975 | 90809.00 | 45.9792 | 96.34 |
| Total | 1988 | 94239.74 | 47.7269 | 100.00 |

**Figure 3.**
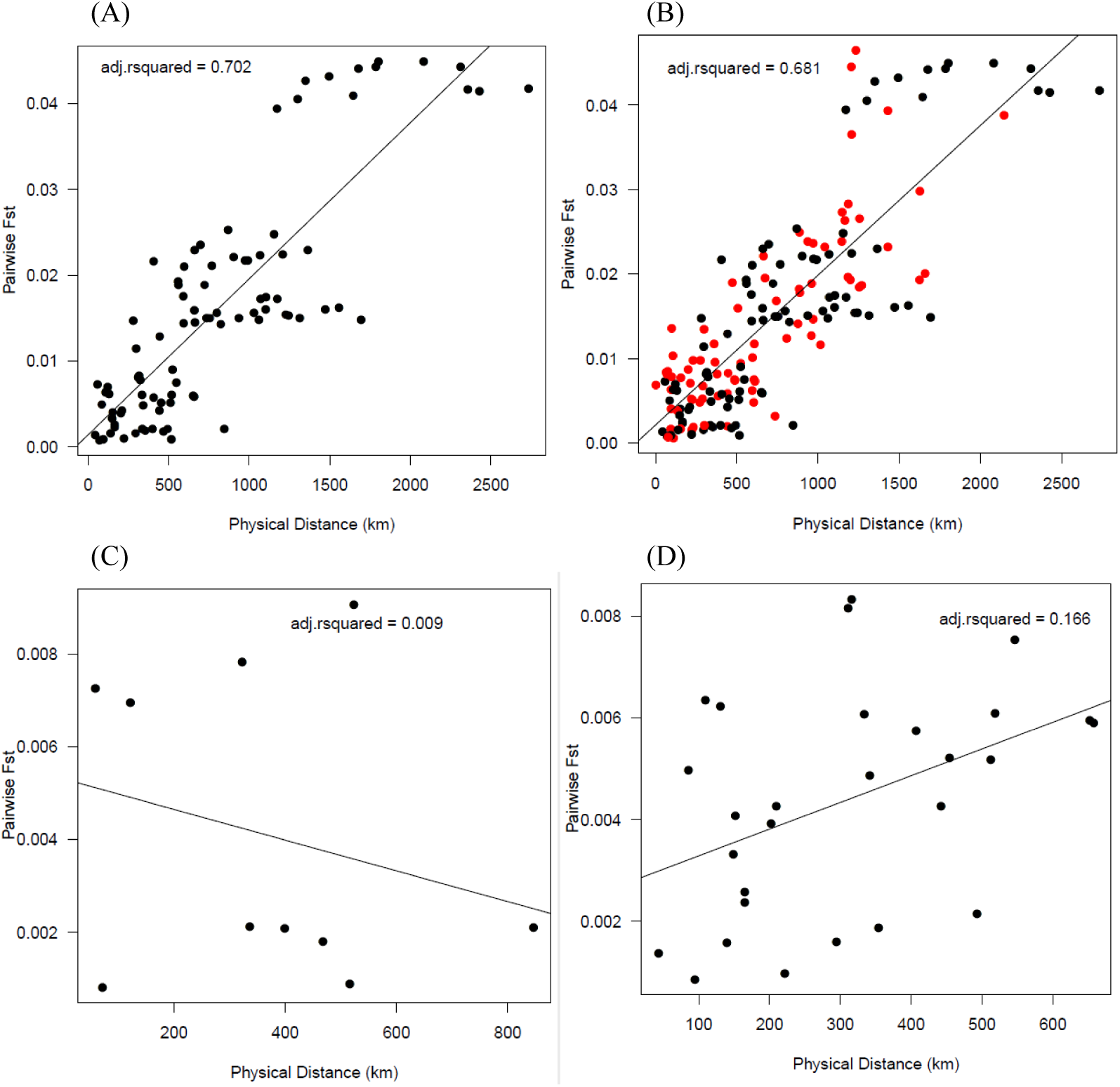
Comparison of physical distance by genetic distance, where each dot represents a comparison of a pair of locations. This indicates that while the populations indicate trends towards isolation-by-distance (A), when looking at only the southern region (C) or the northern region (D) alone, there is no evidence for within-region IBD. Therefore the data is better explained by the hierarchical island model. When including the populations with between 20-35 individuals (B, red dots), the adjusted r-squared value is slightly lower.

### Annual variance in allele frequencies

Applying the minimum sample size threshold of 35, several populations had sufficient sample sizes to split into different years, maintaining the n = 35 threshold for each population-year combination. Populations with multiple year groups included Bella Coola River, Kingcome River, Skeena River, Klamath River, and Fraser River (Figure S5 and Figure S6). For these populations, there was often close clustering of the different collection years, but not always. For example, close clustering occurred for Skeena River 2010 and 2013 (but not 2001), Klamath River 2013 and 2014, Kingcome River 2002 and 2012, Bella Coola River 1998 and 2017 (but not 2013 or 2018). The Fraser River 2014 collection clustered away from the Columbia River collections, but both the 2019 and especially the 2018 collections were more similar to the Columbia River populations, although bootstrap support values for these positions were low. Further demonstration of this by F_ST_ indicates that Bella Coola River 1998 and 2017 collections, Klamath River 2013 and 2014 collections, and Skeena River 2001 vs. 2010 and 2013 were not significantly different from zero (Table 4). The annual variation was highest in the Fraser, with F_ST_ 95% CI ranging from a lower limit of 0.0032 to an upper limit of 0.0106, with the largest difference between Fraser 2014 and 2018.

**Table 4.** Pairwise genetic differentiation between population-year collections using the SNP panel as shown by Weir-Cockerham F_ST_ 95% confidence limits (lower limits in the bottom half, upper limits in the upper half). Shading is used to show increasing values. Only population-year collections with more than 35 individuals are shown. Negative values in the lower limit were replaced by zero and the comparison was considered to not be significantly different.

|  | BEL-1998 | BEL-2017 | BEL-2018 | FRA-2014 | FRA-2018 | FRA-2019 | KIN-2002 | KIN-2012 | KLA-2013 | KLA-2014 | SKE-2001 | SKE-2010 | SKE-2013 |
| --- | --- | --- | --- | --- | --- | --- | --- | --- | --- | --- | --- | --- | --- |
| BEL-1998 | - | 0.0031 | 0.0041 | 0.0227 | 0.0172 | 0.0186 | 0.0052 | 0.0061 | 0.0205 | 0.0219 | 0.0028 | 0.0061 | 0.0065 |
| BEL-2017 | 0 | - | 0.0053 | 0.0276 | 0.0208 | 0.0218 | 0.0071 | 0.0087 | 0.0241 | 0.0272 | 0.0024 | 0.0070 | 0.0060 |
| BEL-2018 | 0.0009 | 0.0011 | - | 0.0250 | 0.0222 | 0.0216 | 0.0076 | 0.0070 | 0.0235 | 0.0264 | 0.0039 | 0.0073 | 0.0068 |
| FRA-2014 | 0.0129 | 0.0160 | 0.0145 | - | 0.0106 | 0.0099 | 0.0226 | 0.0258 | 0.0145 | 0.0135 | 0.0227 | 0.0272 | 0.0313 |
| FRA-2018 | 0.0096 | 0.0112 | 0.0135 | 0.0052 | - | 0.0066 | 0.0135 | 0.0181 | 0.0031 | 0.0035 | 0.0161 | 0.0216 | 0.0213 |
| FRA-2019 | 0.0104 | 0.0121 | 0.0125 | 0.0047 | 0.0032 | - | 0.0208 | 0.0197 | 0.0092 | 0.0107 | 0.0190 | 0.0248 | 0.0285 |
| KIN-2002 | 0.0017 | 0.0023 | 0.0032 | 0.0134 | 0.0067 | 0.0117 | - | 0.0056 | 0.0179 | 0.0185 | 0.0048 | 0.0086 | 0.0064 |
| KIN-2012 | 0.0019 | 0.0040 | 0.0031 | 0.0165 | 0.0097 | 0.0114 | 0.0012 | - | 0.0240 | 0.0241 | 0.0076 | 0.0121 | 0.0099 |
| KLA-2013 | 0.0117 | 0.0131 | 0.0141 | 0.0068 | 0.0002 | 0.0038 | 0.0092 | 0.0131 | - | 0.0024 | 0.0181 | 0.0217 | 0.0231 |
| KLA-2014 | 0.0120 | 0.0146 | 0.0147 | 0.0068 | 0.0005 | 0.0046 | 0.0093 | 0.0139 | 0 | - | 0.0205 | 0.0245 | 0.0262 |
| SKE-2001 | 0 | 0 | 0 | 0.0116 | 0.0064 | 0.0093 | 0 | 0.0019 | 0.0076 | 0.0087 | - | 0.0028 | 0.0042 |
| SKE-2010 | 0.0019 | 0.0010 | 0.0032 | 0.0157 | 0.0120 | 0.0130 | 0.0037 | 0.0063 | 0.0112 | 0.0128 | 0 | - | 0.0036 |
| SKE-2013 | 0.0022 | 0.0014 | 0.0022 | 0.0189 | 0.0115 | 0.0153 | 0.0025 | 0.0047 | 0.0122 | 0.0138 | 0 | 0.0011 | - |

BEL=Bella Coola; COL=Columbia; COW=Cowlitz; FRA=Fraser; KEM=Kemano; KIN=Kingcome; KLA=Klamath; KLI=Klinaklini; NAS=Nass; SAN=Sandy; SKE=Skeena; TWE=Twentymile; UNU=Unuk; WAN=Wannock

### Inter-individual relatedness

Relatedness of individuals within a collection was calculated using both the *wang* and *ritland* estimators (Figure S7). Consistencies were noted between the estimators finding numerous outlier related individuals for the Fraser River, Kingcome River, and Wannock River collections. The *ritland* estimator found highly related individuals within the Twentymile River collection (Figure S7A), but this was not consistent with the *wang* statistic, which showed reduced relatedness for this collection (Figure S7B). Notably, the Ritland statistic is more impacted by siblings in the data (Wang 2002).

### Comparison with microsatellite data

The microsatellite data were obtained from the most recent database that was originally analyzed in Beacham et al. (2005), with augmentation of several stocks including the newly genotyped Klamath River samples. In total, 13 populations were retained that were common with the SNP baseline and that had greater than 35 individuals per population (Figure S8). On average for the microsatellite data, there were 269 individuals per population (sd = 222; min = 69; max = 736).

The general trend of the data was similar between the SNP and microsatellite results, with a large divide between the populations to the south of the Fraser River, inclusive, and the populations to the north of the Fraser River, with Twentymile River as an outgroup (Figure S9). For the microsatellite data, Unuk River was also identified to be an outgroup to the rest of the data. The bootstrap support for the northern and southern general groupings using SNPs (99.99% and 100%, respectively) was higher than that for the microsatellite data (87.92% and 85.81%, respectively). The grouping of the Central Coast and Johnstone Strait (CC-JS) was much less apparent in the microsatellite data, and the Fraser River grouped closely with the Columbia River and south populations in the microsatellite data as well. Interestingly, in both the microsatellite and SNP data, Cowlitz River, Klamath River, and Fraser River grouped together more than any of these grouped with the Columbia River collection (2000). Overall, bootstrap support was higher in the SNP data than the microsatellite, where the SNP data bootstrap values were on average 86.17% (n = 11 values) whereas for the microsatellite data the bootstrap values were on average 64.02% (n = 10 values).

The microsatellite data also showed evidence for IBD across regions (hierarchical island model), with in general a lower overall range of F_ST_ values (Figure S10; F_ST_ range: 0 - ∼0.012) than that observed in the SNP data (Figure 3; F_ST_ range = 0 – ∼0.045), although this was expected due to the different technologies. There was also less of a gap between the regions in the microsatellite data, most notably with the largest distance populations (i.e., Gulf of Alaska vs. southern populations). Further, the trend was less linear for the microsatellite data (microsatellite adj. R^2^ = 0.48; SNP adj. R^2^ = 0.71). In the microsatellite data, some population comparisons were not significantly different from zero (F_ST_ = 0; Table S2). Similarly, the AMOVA for microsatellite data put into regions as determined in the SNP data and geographically (Table S3) shows that only 0.66% of the variation exists between repunits relative to the 2.13% explained in the SNP data. The microsatellite data showed 0.08% of the variation in the data existed between samples within repunit, and 99.26% remaining within samples.

## DISCUSSION

Improved genetic techniques may advance our understanding of the ecologically and culturally important eulachon, for example to address questions about reasons for declines in some populations and healthy returns in others that are in nearby rivers. These techniques may further our understanding of eulachon in the ocean, including their distribution, how stocks mix at-sea, what populations end up in by-catch and where this occurs, and what populations are being characterized in scientific surveys. Here we present a SNP amplicon panel and improved eulachon baseline (14 populations, 1,989 individuals) using 521 differentiating markers sourced from a RADseq study (Candy et al. 2015) combined with a contig-level genome assembly (Sutherland et al. in prep).

The newly developed panel outperforms the existing microsatellite panel (Beacham et al. 2005; Kaukinen et al. 2004) as demonstrated by levels of bootstrap support in dendrograms, improved clustering of populations by geographic region, improved clustering with fewer samples included in the baseline, increased adherence to the expected isolation-by-distance across regions model (hierarchical island model), and increased genetic variance captured within and between groupings (i.e., repunits). Putative adaptive markers in the panel have been found to provide higher levels of differentiation compared to the neutral markers and should provide a better understanding of different selective pressures occurring over the eulachon range (Candy et al. 2015). The new SNP panel is commercially available (Thermo Fisher) and primer sequences are provided herein (Additional File S1).

With the SNP marker panel, three main, large-scale groupings are observed, with some sub-structure within each. This includes the Gulf of Alaska (GoA), southeast Alaska and northern BC, and southern BC through the contiguous US. The northern grouping may yield further subdivision into the Southeast Alaska/North Coast (SEAK-NC) and Central Coast/Johnstone Strait (CC-JS) reporting units, although the true separation of these populations requires additional study through simulations. The southern grouping may yield further subdivision into the Fraser River (FR) and the contiguous US Pacific Northwest (PNW), which also requires further study. Evaluating the effectiveness of more granular resolution within the three identified groups is an important next step for this work, and will be useful to combine both simulated data with empirical data to determine the optimal resolution that can be achieved.

### Regional grouping, isolation-by-distance, and annual variation

Although in general, a hierarchical island model with IBD between regions, but not within regions, explains the variation in the dataset. Within-region IBD was also not identified in a recent study of eulachon in Alaska, where the data were more fit to a hierarchical island model rather than IBD within regions, which was explained as potentially due to large-scale oceanic currents and larval dispersal (Flannery et al. 2013). In the present study, Bella Coola, Klinaklini and Kingcome River populations grouped closely, but the Wannock and Kemano River populations from the similar region grouped separately. The cause of the close grouping of these two separate groups is unknown, but could reflect genetic variation associated with run timing (Table S4) or aspects of habitat differences in the area (e.g., local hydrology). However, the genetic differentiation across these groupings is still low.

For eulachon, run timing may depend on both water temperature and river discharge rate in local river basins (Langer et al. 1977; Ricker et al. 1954; Smith and Saalfeld 1955). Run timing variation can indicate the potential for local adaptation (Beacham *et al*. 2005). Run timing variation occurs across different rivers, generally but not always along a latitudinal gradient (Hay and McCarter 2000; Moody and Pitcher 2010). Peak run timing of eulachon (Table S4) ranges from February in the south (e.g., Columbia River; WDFW & ODFW 2005), March in Central and North coasts (e.g., Kemano, Bella Coola, Skeena, Nass rivers; Moody 2008) or April (e.g., Kingcome and Klinaklini; Southeast Alaska; ADF&G 2008; Moody 2008), and May in Alaska (e.g., Central and Western Alaska; Moody 2008). In contrast to the latitudinal trend, Fraser River (DFO 2020), Sandy River, and Klamath River (Larson and Belchik 1998) have run timing in late March or early April (reviewed in Moody 2008). However, long-term trends toward earlier return timing of eulachon have been noted in several rivers (COSEWIC 2011; Gustafson et al., in prep; Moody and Pitcher 2010), and these trends are likely associated with increasing river temperatures or changes in peak river flows. When environmental conditions are different among locations, and selection acts upon adaptive variants fit to these conditions, local adaptation is possible if sufficient isolation among populations occurs. Low genetic differentiation between populations suggests low drift and/or high gene flow, which reduces but does not preclude the potential for local adaptation. Whether the close genetic similarity observed for Wannock and Kemano river populations and similarly for Bella Coola, Kingcome, and Klinaklini rivers, is due to local environmental differences at each location (e.g., river hydrology) is an interesting avenue for future investigation; additional samples from each of the two different groupings should provide more insight as to whether this trend remains.

The present study found consistent grouping of populations separated by year of collection, such as the Klamath River and Kingcome River, for some but not all collections of Skeena River and Bella Coola River, and slightly higher differentiation across years for the Fraser River. Nonetheless, for the most part, groupings stayed within their putative reporting unit, and always within their larger regions (i.e., the three main groupings). This is consistent with other studies finding a greater effect of geography than temporal variation (Beacham et al. 2005). Sampling multiple years is a useful method of reducing the variance inherent in collections across years (Waples 1998), and has been highlighted as a valuable step to evaluate potential for mixed-stock analysis accuracy (Flannery et al. 2013). In addition, with climate change scenarios and expected changes in the distribution of species, it will be informative to continue collecting baseline samples in future years to ensure trends remain consistent. It is interesting to note that there may be multiple runs per river, having peak spawning time at different dates, as has been observed in the Nass River, with an initial run arriving in early to mid March, and a second run arriving in early April (COSEWIC 2013; Langer et al. 1977; Noble et al. 2012), as well as the Kingcome River (Gustafson et al. 2010), the Fraser River (LFFA 2015), the Elwha River (Gustafson 2016), and others throughout the range. If this hidden variation within a river is not included, for example in metadata of collections, it could lead to variance occurring across sampling years if there is only a single sampling event per year.

### Genetics and current management groupings

Gustafson et al. (2012) identified one Distinct Population Segment (DPS) of eulachon in the California Current. DPSs to the north of the Skeena River were not identified, as the Status Review teams’ mandate was to identify a DPS of eulachon that contained the petitioned populations of eulachon from the states of California, Oregon, and Washington, and not to identify DPSs coastwide. Gustafson et al. (2012) suggest the strong ecological and environmental break that occurs between the Alaska and California Currents provide support for discreteness of eulachon within the California Current.

The three Designatable Units (DUs) within Canada (COSEWIC 2011) correspond well with genetic structure observed here, although it is not entirely clear where the separation point exists between the Nass/Skeena DU and the Central Coast, either around Bella Coola or Johnstone Strait. The Nass and Skeena River populations are very similar genetically to the Kemano and Wannock River populations, suggesting these should be collectively considered as one grouping. However, the Nass and Skeena River populations are also very similar genetically to the Bella Coola River population Whether the Kingcome, Bella Coola, and Klinaklini rivers are distinct enough from these other Central and North Coast populations for sufficient resolution in mixed-stock analysis is an important next step to this work. The present results are in agreement with previous results suggesting high gene flow between the Nass River and Skeena River populations (see COSEWIC 2013). Additional baseline samples in future years should continue to help resolve the groupings and monitor for any changes. These will be continued to be added to the existing baseline, as multiple year collections are known to reduce the potential for single year sampling biases.

## CONCLUSIONS

Based on genetic evidence, the current baseline with populations from Klamath River in California through coastal British Columbia and north to Twentymile River in Alaska is tentatively grouped into five reporting units. The current SNP panel outperformed the microsatellite panel and will be applied to mixed-stock analysis. Low genetic differentiation was observed overall compared to other anadromous species such as salmonids, and a hierarchical island model best explained the structure observed here. Annual differences in collections were characterized, and although in general collections clustered together regardless of year, some variance existed, indicating the value of having these multiple year collections in the data. Several clustering trends remain unexplained, but may be related to run timing or hydrological differences of the rivers. Although improvements are expected as the baseline continues to grow, the current baseline is a foundation that will be used for subsequent mixed-stock analysis. Important next steps will involve simulating and testing mixed-stock samples to determine the reliability of the five reporting units proposed here from the population structure analysis.

## ACKNOWLEDGEMENTS

This work was supported by funds from Fisheries and Oceans Canada (DFO) Rotational Survey Fund, as well as the Species At Risk program (SARA). We thank First Nations and Fisheries and Oceans Canada staff for the eulachon collections along the coast of British Columbia, staff from the Washington Department of Fisheries (DFW) for collections from Washington, and staff from Alaska Department of Fish & Game, US Fish & Wildlife Service, and the University of Alaska Fairbanks for collections from Alaska. Thanks to the following people and groups for samples collected since 2015: Bella Coola River samples were provided by the Nuxalk Nation (Jason Moody and Megan Moody); Fraser River samples were provided by the Lower Fraser Fisheries Alliance (LFFA); Klamath River samples were provided by the Yurok Tribal Fisheries Program, Klamath River Division (Barry McCovey Jr. and Michael Belchik); Elwha River samples were provided by the Lower Elwha Klallam Tribe (Mike McHenry); Skeena River 2019 samples were provided by DFO (Vanessa Hodes and Lindsay Dealy); Sandy River samples were provided by Maureen Small of DFW; and Nass and Skeena River (2013), Unuk River, and Wannock River samples were provided by DFO (Matt Thompson). Thanks to Ashtin Duguid (DFO) and Ana Ramon-Laca (NMFS) for help with additional DNA extractions. Thanks to Haktan Suren and Prasad Siddavatam and the rest of the Thermo Fisher Scientific team for development of commercial primers for eulachon. Thanks to Alejandro Frid and Megan Moody for comments on an earlier version of the manuscript.

## DATA ACCESSIBILITY

The raw data for this analysis is available on the NCBI Short Read Archive (SRA) under BioProject Accession PRJNA635905 within BioSamples SAMN15057088-SAMN15060309. The original RAD-seq data are available within Candy et al. (2015). The eulachon ampliseq panel can be ordered using the following catalogue number from Thermo Fisher: SKU A44467 AgriSeq Custom Panel –DFO_EULACHON20180627. Additional Files including the primers for the eulachon panel and the hotspot file outlining variants and positions can be found on FigShare: https://doi.org/10.6084/m9.figshare.12922538.v2 The analytical pipelines applied in this work are all available on GitHub, including: Extend RAD marker using genome: https://github.com/bensutherland/fasta_SNP_extraction Analyze population genetic data: https://github.com/bensutherland/simple_pop_stats

## SUPPLEMENTAL INFORMATION

**Additional File S1**. Primers for the eulachon panel.

**Additional File S2**. Hotspot file outlining variants and positions.

**Figure S1**. Amplicon panel observed heterozygosity per marker.

**Figure S2**. Amplicon panel dendrogram showing genetic similarity among populations in the baseline including all populations that have at least 20 individuals.

**Figure S3**. (A) Principle Components Analysis of the SNP baseline showing PC1 and PC2; and (B) eigenvalues of the different PCs.

**Figure S4**. Principle component loading values of each marker for the top three PCs.

**Figure S5**. Sample sizes in the amplicon panel baseline when separating by location and year.

**Figure S6**. Amplicon panel genetic dendrogram when considering collections separated by year.

**Figure S7**. Inter-individual relatedness within each population estimated from the data in the amplicon baseline

**Figure S8**. The microsatellite baseline sample size per collection

**Figure S9**. The microsatellite baseline dendrogram showing genetic similarity among collections. Colours represent repunits as determined by the amplicon panel.

**Figure S10**. Physical by genetic distance comparison in the microsatellite panel

**Table S1**. Amplicon panel pairwise genetic differentiation estimates.

**Table S2**. Microsatellite panel pairwise genetic differentiation estimates.

**Table S3**. Microsatellite panel variance sources as determined by Analysis of Molecular Variance (AMOVA).

**Table S4**. Run timing differences for eulachon populations relevant to the study.

