## Supplemental Information for "Population structure of eulachon *Thaleichthys pacificus* from Northern California to Alaska using single nucleotide polymorphisms from direct amplicon sequencing"

Supplemental results and information for the article “Population structure of eulachon *Thaleichthys pacificus* from Northern California to Alaska using single nucleotide polymorphisms from direct amplicon sequencing”.

**SUPPLEMENTAL FIGURES**

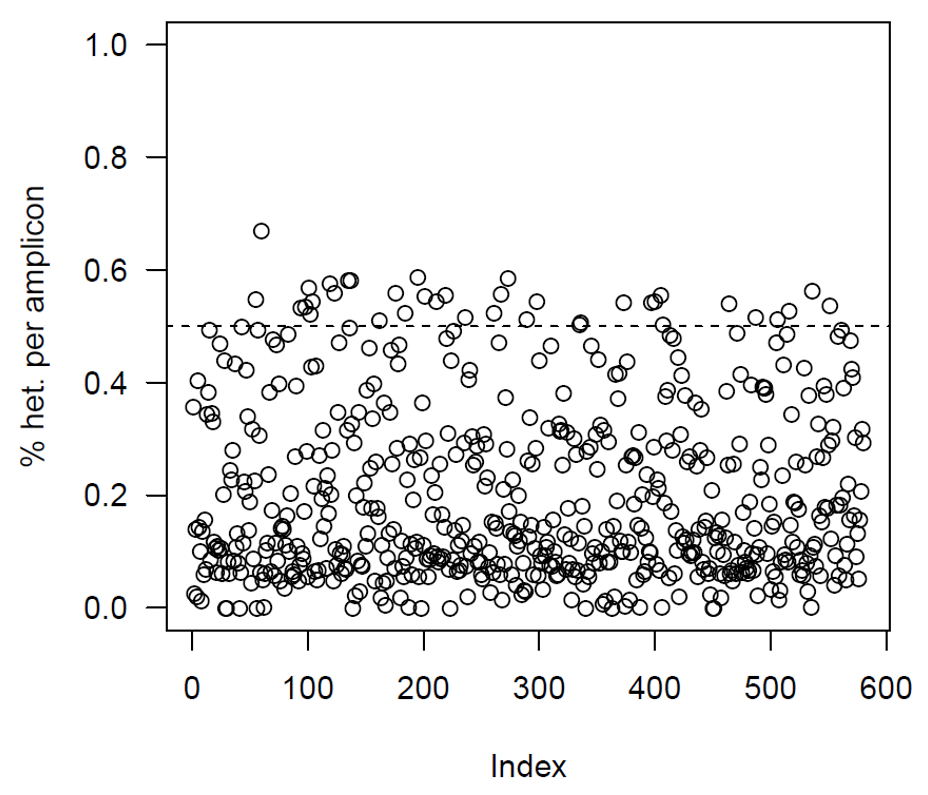

**Figure S1.** Amplicon panel observed heterozygosity per marker (single SNP per amplicon) across all samples included in the study (including all collections with more than 20 individuals). A total of 25 markers showed heterozygosity greater than 0.5, and 14 markers had heterozygosity less than 0.01.

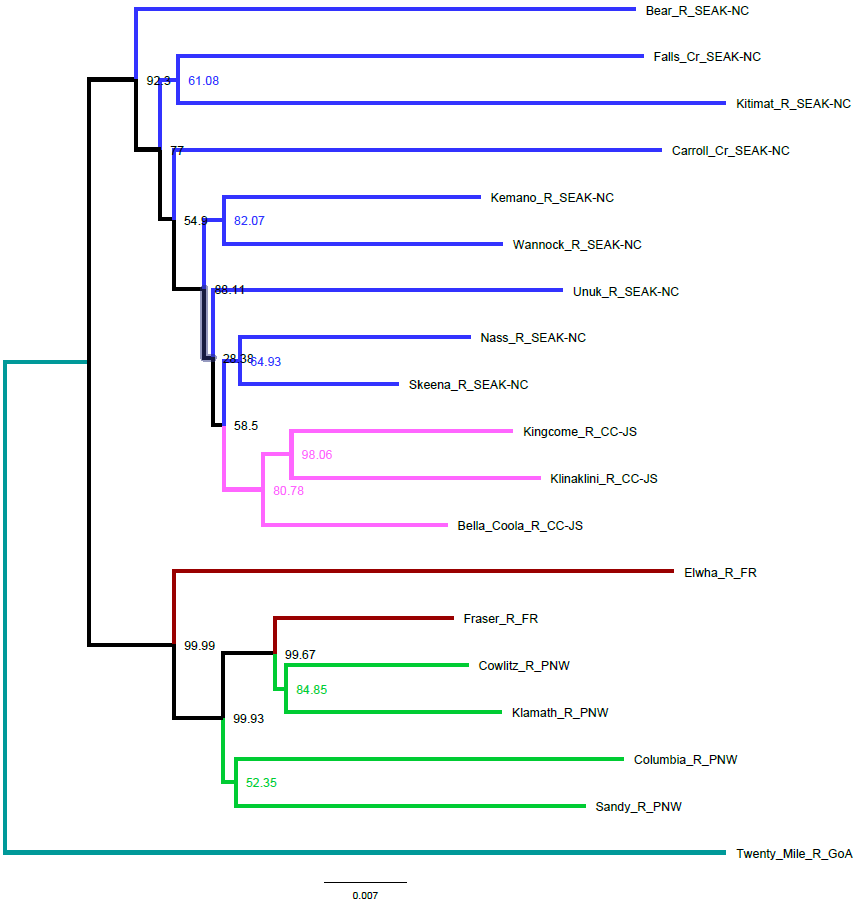

**Figure S2**. Amplicon panel dendrogram showing genetic similarity among populations in the baseline including all populations that have at least 20 individuals (i.e., not final filtered baseline). Poor resolving power was observed for collections with fewer than 35 individuals, as observed by lack of clustering within geographic regions (e.g., Carroll Creek, Kitimat River, Falls Creek, and Bear River).

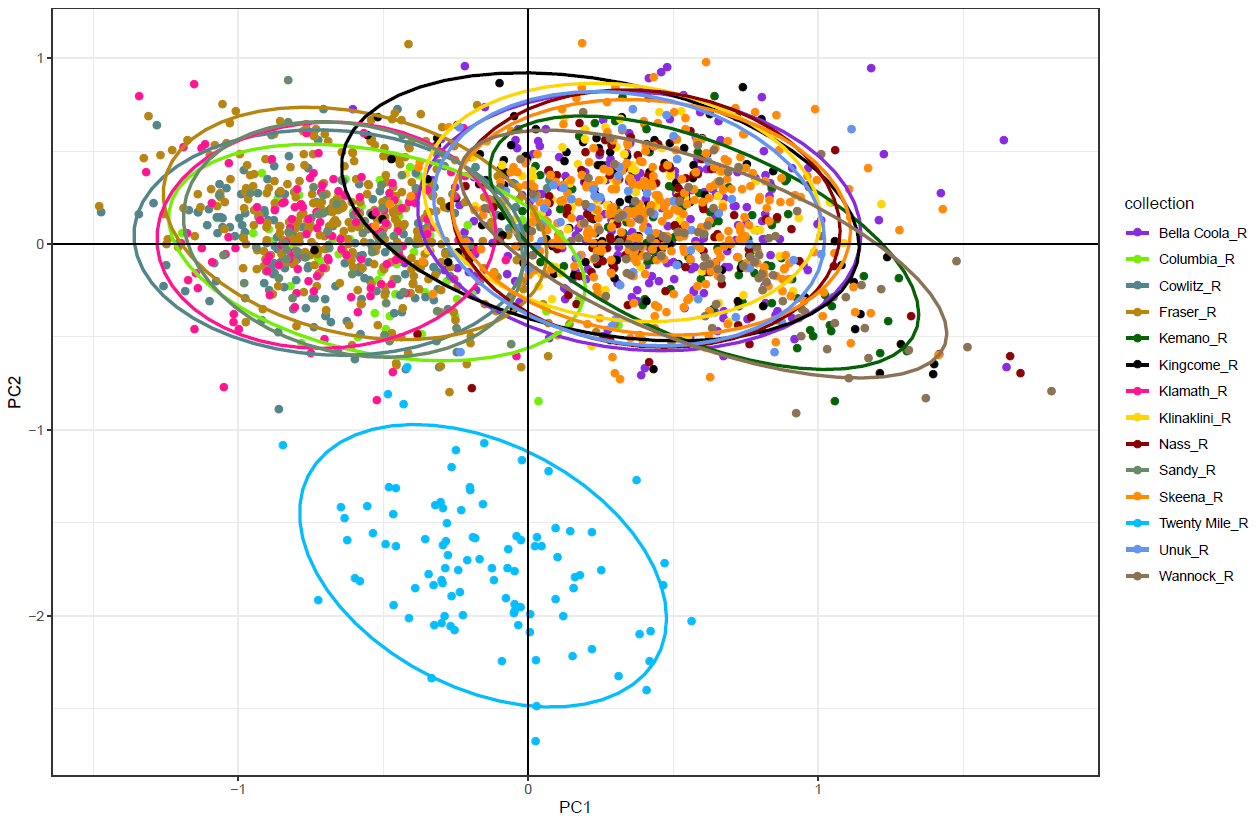

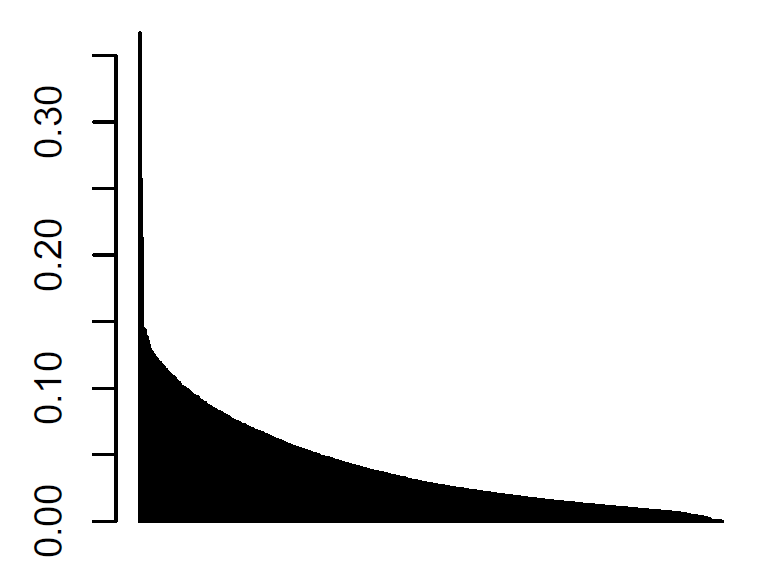

**Figure S3**. Using the amplicon panel, (A) a PCA indicates a strong separation along PC2 between the Twentymile River (AK) sample and the rest of the samples, as well as two main groupings (south in negative PC1; north in positive PC1). The ellipses show 95% confidence intervals around the samples from each group. (B) PCA eigenvalues indicates the importance of retaining the first three PCs.

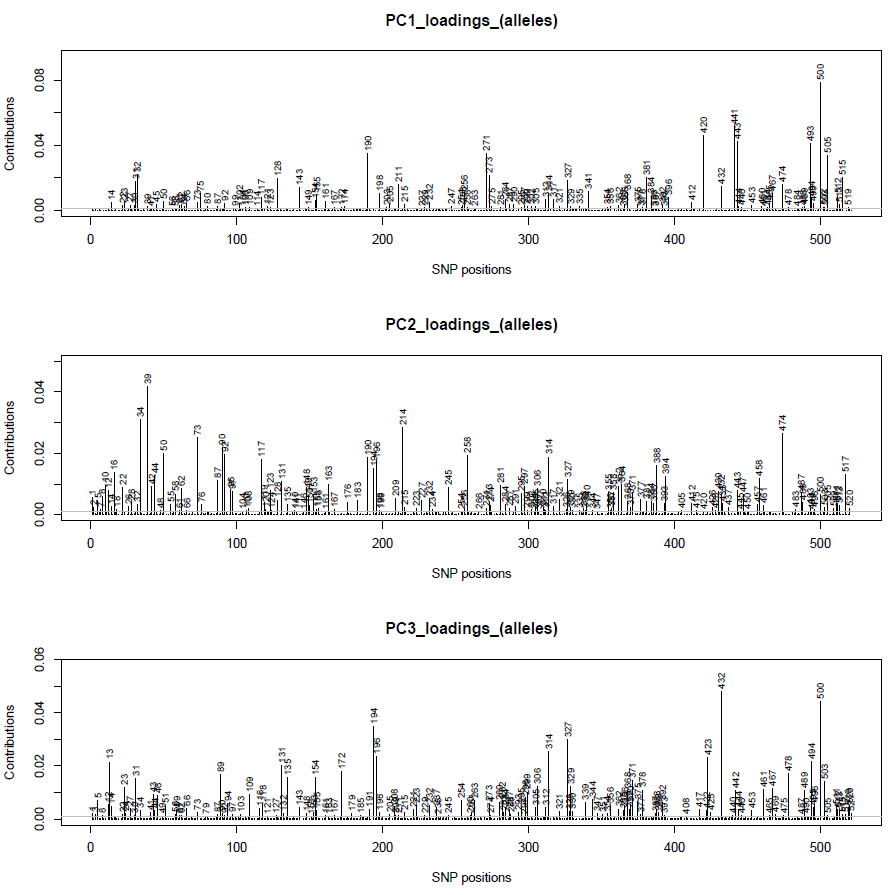

**Figure S4**. Loadings of markers by index and their contributions to separations along PC1, PC2 and PC3 from Figure S3 indicates that many markers are involved in separating PCs.

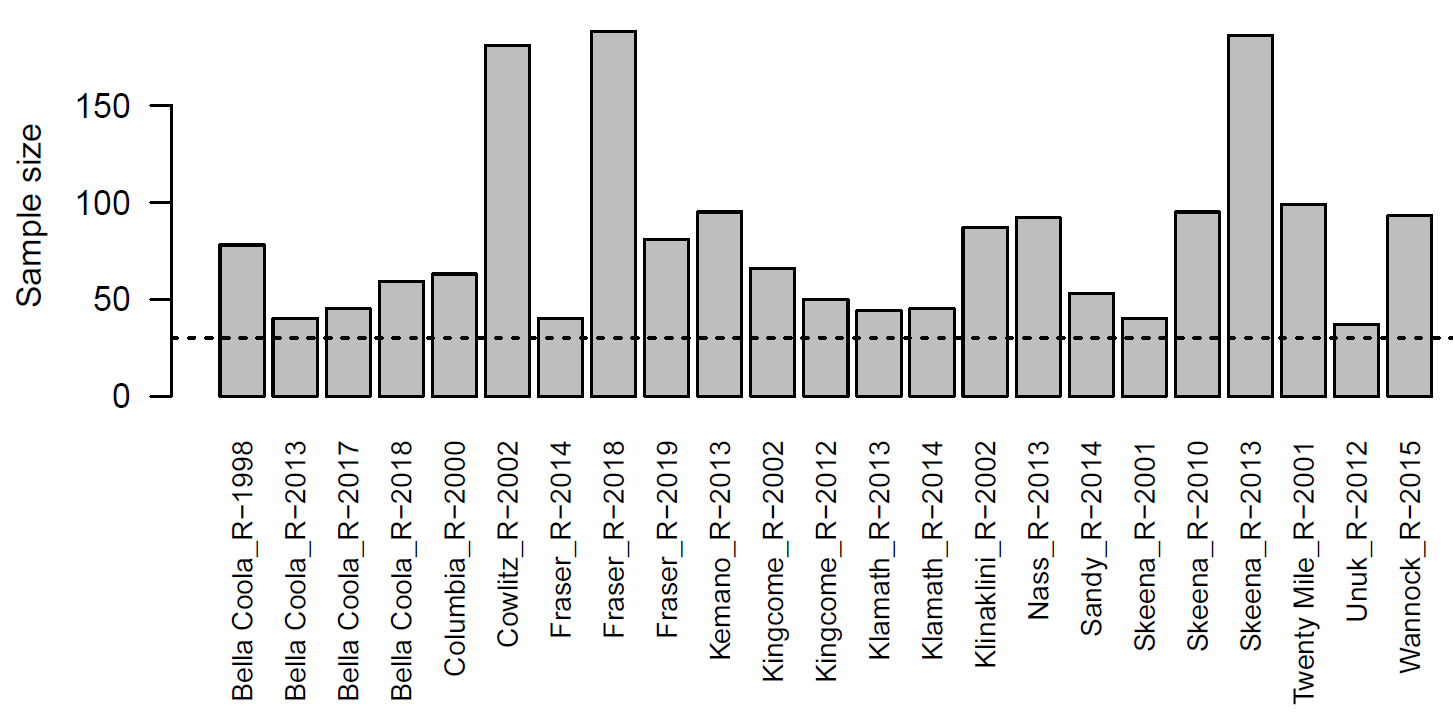

**Figure S5**. Sample sizes in the amplicon panel baseline when separating by location and year for all populations with more than 35 individuals per collection.

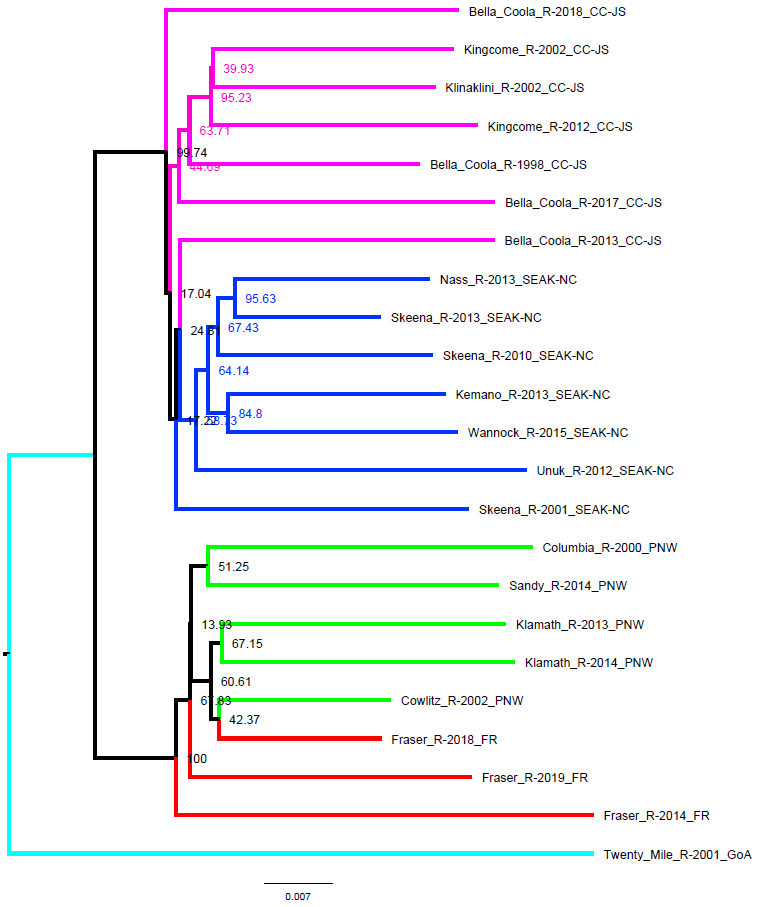

**Figure S6**. Amplicon panel genetic dendrogram when considering collections separated by year, only considering those year-collection combinations with at least 35 individuals.

**(A)**

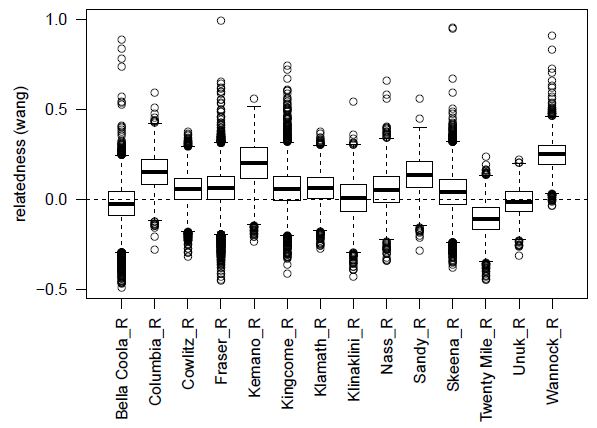

**(B)**

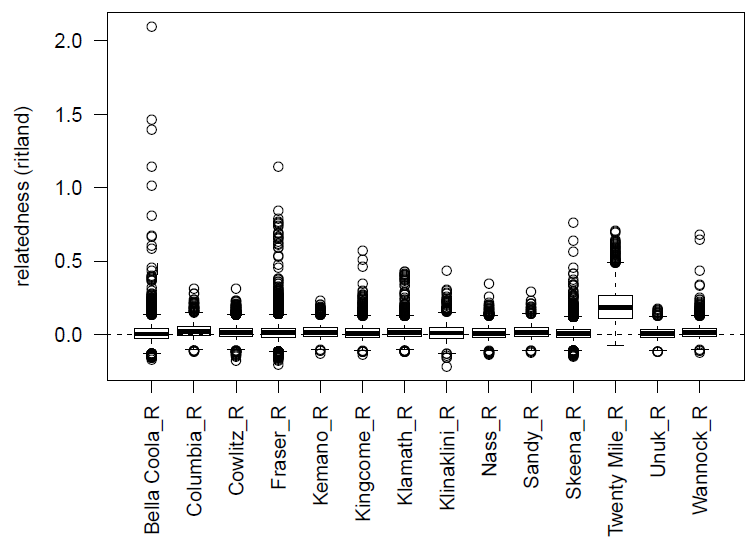

**Figure S7.** Inter-individual relatedness within each population estimated from the data in the amplicon baseline using either the (A) Wang estimator; or (B) Ritland estimator.

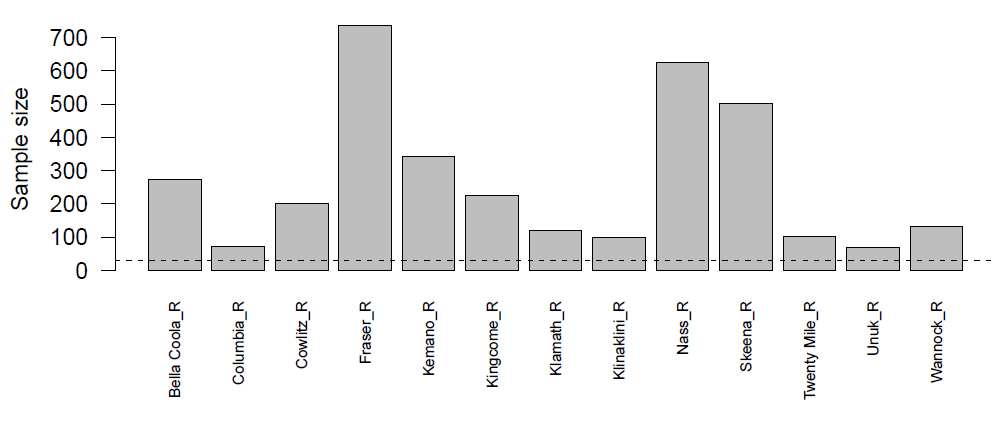

**Figure S8.** The microsatellite baseline sample size per collection.

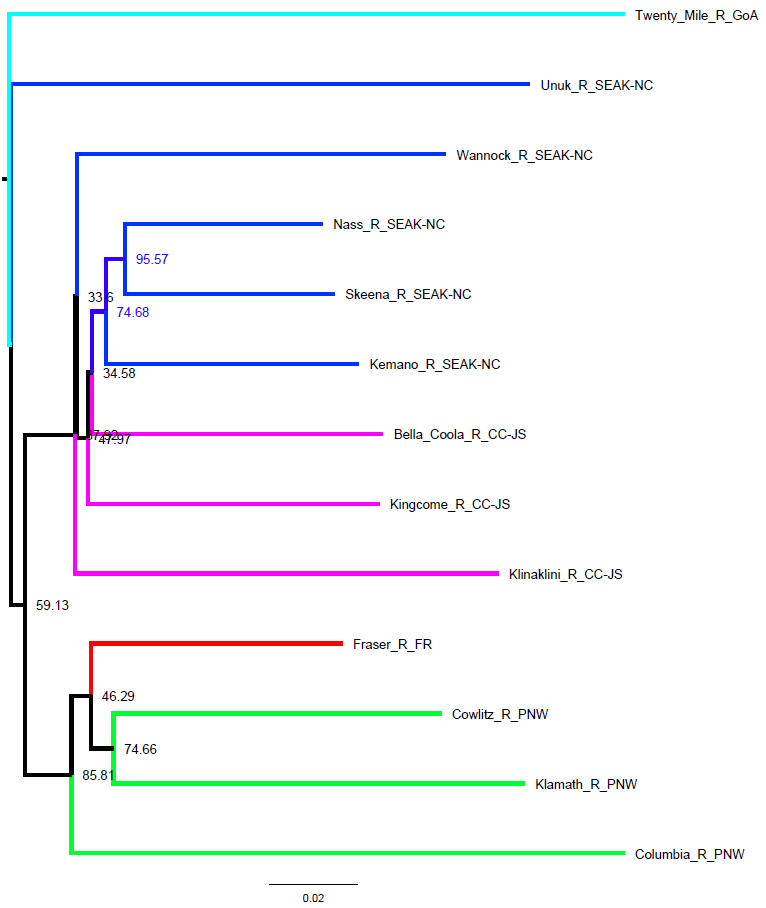

**Figure S9.** The microsatellite baseline dendrogram showing genetic similarity among collections. Colours represent repunits as determined by the amplicon panel.

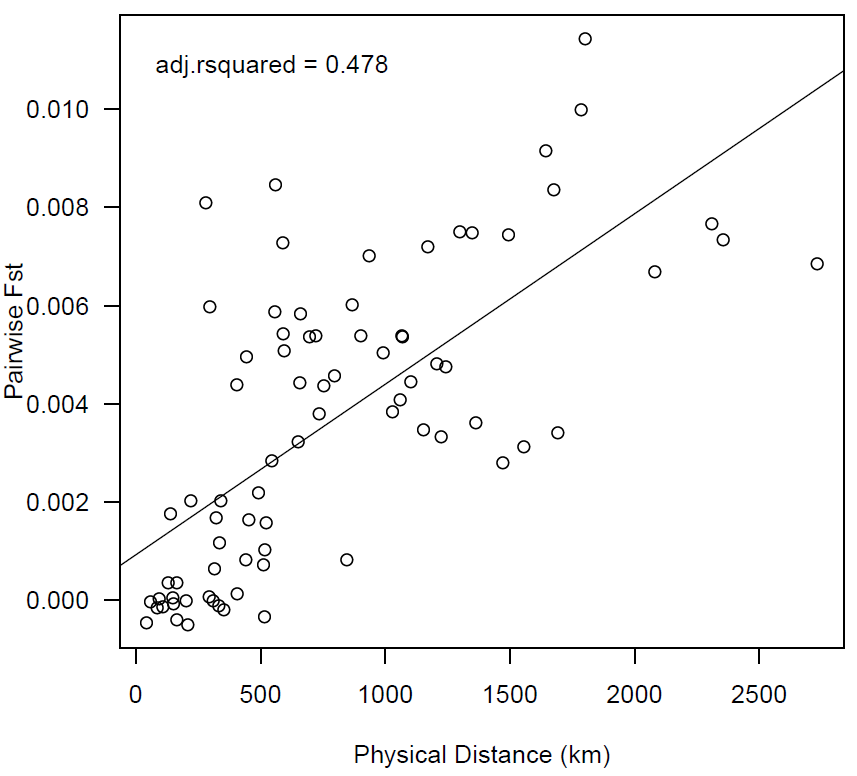

**Figure S10**. The microsatellite baseline showing a similar relationship between physical and genetic distance (Weir-Cockerham F_ST_) to the SNP panel (hierarchical island mode), although with a reduced adjusted R-squared value for model fit.

**SUPPLEMENTAL TABLES**

**Table S1.** Amplicon panel pairwise genetic differentiation estimates (Weir-Cockerham F_ST_) between all populations in the filtered baseline. Shading is used to show increasing F_ST_ values.

|  | BEL | COL | COW | FRA | KEM | KIN | KLA | KLI | NAS | SAN | SKE | TWE | UNU | WAN |
| --- | --- | --- | --- | --- | --- | --- | --- | --- | --- | --- | --- | --- | --- | --- |
| BEL | NA | 0.0189 | 0.0150 | 0.0129 | 0.0041 | 0.0024 | 0.0155 | 0.0026 | 0.0019 | 0.0143 | 0.0016 | 0.0410 | 0.0021 | 0.0050 |
| COL |  | NA | 0.0073 | 0.0079 | 0.0254 | 0.0193 | 0.0091 | 0.0189 | 0.0224 | 0.0070 | 0.0217 | 0.0443 | 0.0225 | 0.0230 |
| COW |  |  | NA | 0.0021 | 0.0222 | 0.0144 | 0.0009 | 0.0176 | 0.0175 | 0.0008 | 0.0157 | 0.0417 | 0.0154 | 0.0235 |
| FRA |  |  |  | NA | 0.0211 | 0.0114 | 0.0021 | 0.0148 | 0.0157 | 0.0021 | 0.0150 | 0.0449 | 0.0151 | 0.0217 |
| KEM |  |  |  |  | NA | 0.0082 | 0.0230 | 0.0083 | 0.0039 | 0.0218 | 0.0033 | 0.0433 | 0.0049 | 0.0043 |
| KIN |  |  |  |  |  | NA | 0.0148 | 0.0014 | 0.0052 | 0.0146 | 0.0043 | 0.0443 | 0.0060 | 0.0064 |
| KLA |  |  |  |  |  |  | NA | 0.0173 | 0.0163 | 0.0018 | 0.0161 | 0.0418 | 0.0149 | 0.0248 |
| KLI |  |  |  |  |  |  |  | NA | 0.0061 | 0.0160 | 0.0052 | 0.0450 | 0.0059 | 0.0062 |
| NAS |  |  |  |  |  |  |  |  | NA | 0.0172 | 0.0009 | 0.0406 | 0.0016 | 0.0058 |
| SAN |  |  |  |  |  |  |  |  |  | NA | 0.0161 | 0.0415 | 0.0151 | 0.0212 |
| SKE |  |  |  |  |  |  |  |  |  |  | NA | 0.0428 | 0.0010 | 0.0061 |
| TWE |  |  |  |  |  |  |  |  |  |  |  | NA | 0.0395 | 0.0442 |
| UNU |  |  |  |  |  |  |  |  |  |  |  |  | NA | 0.0076 |
| WAN |  |  |  |  |  |  |  |  |  |  |  |  |  | NA |

**Table S2.** Microsatellite panel pairwise genetic differentiation estimates (Weir-Cockerham F_ST_) between populations. Shading is used to show increasing F_ST_ values.

|  | BEL | COL | COW | FRA | KEM | KIN | KLA | KLI | NAS | SKE | TWE | UNU | WAN |
| --- | --- | --- | --- | --- | --- | --- | --- | --- | --- | --- | --- | --- | --- |
| BEL | NA | 0.0054 | 0.0044 | 0.0050 | 0 | 0 | 0.0033 | 0.0004 | 0 | 0.0001 | 0.0091 | 0.0022 | 0 |
| COL |  | NA | 0.0000 | 0.0017 | 0.0060 | 0.0059 | 0.0016 | 0.0085 | 0.0054 | 0.0051 | 0.0077 | 0.0048 | 0.0058 |
| COW |  |  | NA | 0.0012 | 0.0054 | 0.0054 | 0 | 0.0073 | 0.0045 | 0.0038 | 0.0073 | 0.0048 | 0.0054 |
| FRA |  |  |  | NA | 0.0051 | 0.0060 | 0.0008 | 0.0081 | 0.0046 | 0.0038 | 0.0067 | 0.0070 | 0.0044 |
| KEM |  |  |  |  | NA | 0.0000 | 0.0036 | 0.0007 | 0.0000 | 0.0000 | 0.0075 | 0.0020 | 0 |
| KIN |  |  |  |  |  | NA | 0.0041 | 0 | 0.0007 | 0.0008 | 0.0100 | 0.0032 | 0 |
| KLA |  |  |  |  |  |  | NA | 0.0054 | 0.0031 | 0.0028 | 0.0068 | 0.0034 | 0.0035 |
| KLI |  |  |  |  |  |  |  | NA | 0.0010 | 0.0016 | 0.0114 | 0.0044 | 0.0004 |
| NAS |  |  |  |  |  |  |  |  | NA | 0.0000 | 0.0075 | 0.0018 | 0.0001 |
| SKE |  |  |  |  |  |  |  |  |  | NA | 0.0075 | 0.0020 | 0 |
| TWE |  |  |  |  |  |  |  |  |  |  | NA | 0.0072 | 0.0084 |
| UNU |  |  |  |  |  |  |  |  |  |  |  | NA | 0.0028 |
| WAN |  |  |  |  |  |  |  |  |  |  |  |  | NA |

**Table S3.** Microsatellite panel variance sources as determined by Analysis of Molecular Variance (AMOVA). The same populations and grouping structure as the amplicon panel were used.

| **Source of variation** | **d.f.** | **Sums of squares** | **Variance components (sigma)** | **Percentage of variation** |
| --- | --- | --- | --- | --- |
| Between repunit | 4 | 113.74 | 0.0361 | 0.66 |
| Between samples within repunit | 8 | 51.34 | 0.0044 | 0.08 |
| Within samples | 3487 | 18924.08 | 5.4270 | 99.26 |
| Total | 3499 | 19089.16 | 5.4675 | 100.00 |

**Table S4.** Run timing differences for eulachon populations relevant to the study. The differences are not always clinal, and although Washington has peak spawning in February, Northern California and Fraser River have peak spawning in late March-early May. The most Northern populations spawn in May.

| **Month of Peak Spawning Abundance** | **Population** |
| --- | --- |
| February | Washington / Oregon (South) ^1,2^  Columbia R (South)^1,2^ |
| March | Kemano R (Central)^1^  Bella Coola R (Central)^1^  Nass R (North)^1^  Skeena R (North)^1^  Sandy R. (South)^3^ |
| March/April | Northern California (South)^4^ |
| April | Fraser River (South)^1^  Kingcome R (Central)^1^  Klinaklini R (Central)^1^  Southeast Alaska (AK)^5^ |
| April/May | Fraser River (South)^6^ |
| May | Central Alaska (AK)^1^  Western Alaska (AK)^1^ |

^1^ Moody, 2008; ^2^ WDFW & ODFW 2005; ^3^ Smith & Saalfeld, 1955; ^4^ Larson & Belchik, 1998; ^5^ ADF&G, 2008; ^6^ DFO 2020; see References for full citations.
